# Loss of the Na^+^/K^+^ cation pump CATP-1 suppresses *nekl*-associated molting defects

**DOI:** 10.1101/2024.03.15.585189

**Authors:** Shaonil Binti, Phil T. Edeen, David S. Fay

## Abstract

The conserved *C. elegans* protein kinases NEKL-2 and NEKL-3 regulate multiple steps of membrane trafficking and are required for larval molting. Through a forward genetic screen we identified a loss-of-function mutation in *catp-1* as a suppressor of molting defects in synthetically lethal *nekl-2; nekl-3* double mutants. *catp-1* is predicted to encode a membrane-associated P4-type ATPase involved in Na^+^–K^+^ exchange. Moreover, a mutation predicted to abolish CATP-1 ion-pump activity also suppressed *nekl-2; nekl-3* mutants. Endogenously tagged CATP-1 was primarily expressed in epidermal (hypodermal) cells within punctate structures located at or near the apical plasma membrane. Through whole genome sequencing, we identified two additional *nekl-2; nekl-3* suppressor strains containing coding-altering mutations in *catp-1* but found that neither mutation, when introduced into *nekl-2; nekl-3* mutants using CRISPR methods, was sufficient to elicit robust suppression of molting defects. Our data also suggested that the two *catp-1* isoforms, *catp-1a* and *catp-1b*, may in some contexts be functionally redundant. On the basis of previously published studies, we tested the hypothesis that loss of *catp-1* may suppress *nekl*-associated defects by inducing partial entry into the dauer pathway. Contrary to expectations, however, we failed to obtain evidence that loss of *catp-1* suppresses *nekl-2; nekl-3* defects through a dauer-associated mechanism or that loss of *catp-1* leads to entry into the pre-dauer L2d stage. As such, loss of *catp-1* may suppress *nekl-* associated molting and membrane trafficking defects by altering electrochemical gradients within membrane-bound compartments.

## INTRODUCTION

Ecdysozoa, animals that undergo molting encompass the major phyla Nematoda (roundworms) and Arthropoda (insects, chelicerates, crustaceans, and myriapods), as well as several smaller phyla including Tardigrada (tardigrades) and Nematomorpha (parasitic horsehair worms) (AGUINALDO *et al*. 1997; EWER 2005; TELFORD *et al*. 2008). Common to ecdysozoans is a multi-layered exoskeleton, the cuticle, which is an apical extracellular matrix secreted primarily by the epidermis. The material composition and mechanical properties of the cuticle can vary widely among different species and life stages. Whereas the cuticle is essential for movement, protection, and other basic life functions, it can pose limitations on developmental processes including animal growth. To accommodate growth and morphological changes, ecdysozoans undergo cycles of molting (ecdysis), whereby a new cuticle is synthesized underneath the old cuticle, which is shed. In the case of the nematode *Caenorhabditis elegans*, the shedding process occurs at the termination of each larval stage (L1, L2, L3, and L4), with the new cuticle allowing for body size expansion when resources are replete, or long-term survival (as dauer larvae; L3d) when conditions are deemed adverse (SINGH AND SULSTON 1978; FIELENBACH AND ANTEBI 2008; LAZETIC AND FAY 2017b).

Progression through molting requires the coordination of several cellular processes including the ordered transcription of hundreds of molting-associated genes; the secretion and construction of a new cuticle from the outside in; and the detachment, degradation, and partial recycling of the old cuticle (JOHNSTONE AND BARRY 1996; PAGE AND JOHNSTONE 2007; LAZETIC AND FAY 2017b; MIAO *et al*. 2020; TSIAIRIS AND GROSSHANS 2021). Correspondingly, mutations that disrupt these processes within the epidermal cells that underly and secrete the cuticle can lead to defective molting and larval arrest. These include mutations affecting several steroid-hormone receptors required for the cyclic transcription of molting genes, mutations in individual cuticle components and apical extracellular matrix regulators, and mutations that affect membrane trafficking including endocytosis and exocytosis (ASAHINA *et al*. 2000; GISSENDANNER AND SLUDER 2000; KOSTROUCHOVA *et al*. 2001; FRAND *et al*. 2005; MONSALVE AND FRAND 2012; LAZETIC AND FAY 2017b; JOHNSON *et al*. 2023). In addition, loss of individual plasma-membrane proteins can lead to molting arrest. One example is LRP-1/Megalin, an epidermal apolipoprotein receptor required for the internalization of environmental sterols via clathrin-mediated endocytosis (YOCHEM *et al*. 1999; KANG *et al*. 2013). Because cholesterol uptake is essential for the synthesis of steroid hormones in *C. elegans*, a failure to internalize LRP-1 may preclude the normal transcriptional activation of molting genes (MERRIS *et al*. 2003; ENTCHEV AND KURZCHALIA 2005; MARTIN *et al*. 2010; LAZETIC AND FAY 2017b; JOSEPH *et al*. 2020; BINTI *et al*. 2022).

Through genetic screens, we previously identified two *C. elegans* NimA-related kinases, NEKL-2 and NEKL-3, as essential for the completion of molting (YOCHEM *et al*. 2015). Specifically, null alleles of *nekl-2* or *nekl-3* lead to penetrant molting defects and larval arrest at the L1/L2 transition, whereas partial loss-of-function alleles cause larvae to arrest at L2/L3. NEKL-2 is most similar to human NEK8 and NEK9, whereas NEKL-3 is orthologous to NEK6 and NEK7.

Consistent with these findings, our genetic screens identified conserved ankyrin-repeat partners of NEKL-2 (MLT-2/ANKS6 and MLT-4/INVS) and NEKL-3 (MLT-3/ANKS3) as required for molting, which we showed are necessary for proper NEKL subcellular localization (LAZETIC AND FAY 2017a). Further studies have indicated roles for NEKL-2 and NEKL-3 at multiple steps of membrane trafficking, including clathrin-mediated endocytosis and endosomal transport, and have implicated human NEK6 and NEK7 in analogous trafficking functions in tissue culture cells (JOSEPH *et al*. 2020; JOSEPH *et al*. 2023).

In previous work, we described a genetic screen to identify suppressors of molting defects in *nekl* mutants (JOSEPH *et al*. 2018). Specifically, we combined weak aphenotypic loss-of-function alleles of *nekl-2* and *nekl-3* to create synthetically lethal *nekl-2(fd81); nekl-3(gk894345)* double mutants, which were used to identify suppressors of *nekl* synthetic lethality. Our screen led to the identification of mutations affecting several core components of membrane trafficking as well as ECM modifiers and cell signaling components (JOSEPH *et al*. 2018; LAZETIC *et al*. 2018; JOSEPH *et al*. 2022). In addition, our studies demonstrated that mutations and environmental conditions that promote induction of the dauer pathway can suppress a subset of *nekl* alleles that arrest at the L2/L3 transition (BINTI *et al*. 2022). Namely, entry into the uncommitted pre-dauer state, L2d, can be sufficient to suppress molting defects and to partially restore the expression of molting-associated genes.

Here we report a novel *nekl*-suppressor mutation affecting CATP-1, a P4-type Na^+^–K^+^ pump. CATP-1 functions to maintain an electrochemical gradient across membranes by exporting three Na^+^ ions and importing two K^+^ ions during each ATP-driven catalytic cycle (KAPLAN 2002; MORTH *et al*. 2011; CLAUSEN *et al*. 2017). Loss of *catp-1* suppresses larval arrest caused by exposure of worms to dimethylphenylpiperazinium (DMPP) (RUAUD AND BESSEREAU 2007), a nicotinic agonist that uncouples the coordinated timing of cell divisions with molting cycles (RUAUD AND BESSEREAU 2006). This study also provided evidence that CATP-1 is expressed and functions in the epidermis, consistent with a possible role in molting. In addition, evidence suggested that CATP-1 may be an effector of the dauer pathway, potentially linking to our finding that dauer induction can partially bypass the requirement for NEKL kinases (BINTI *et al*. 2022). More recently, CATP-1 was shown to function in glial cells that associate with mechanosensory neurons involved in touch sensitivity (JOHNSON *et al*. 2020). Our current studies demonstrate that loss of CATP-1 ion-pump activity can suppress molting defects in *nekl-2(fd81); nekl-3(gk894345)* mutants. In contrast to expectations, however, our data do not support the model that *catp-1*–mediated suppression occurs through induction of the dauer pathway, raising the possibility that changes in ion balance may impact molting through other mechanisms including possible effects on membrane trafficking.

## MATERIALS AND METHODS

### Strains and maintenance

*C. elegans* strains were cultured on nematode growth medium (NGM) spotted with *E. coli* OP50 and they were maintained at 22°C (unless otherwise stated) according to standard protocols (Stiernagle, 2006). All strains used in this study are listed in Supplementary File 1.

### Genome sequencing

Whole genome sequencing (WGS) of strains WY1286 (5× backcross), WY1744 (3× backcross), and WY1768 (1× backcross) was carried out as described (JOSEPH *et al*. 2018). After sequencing of WY1286, the Sibling Subtraction Method (JOSEPH *et al*. 2018)was carried out to reduce the number of candidate causal mutations.

### CRISPR/Cas9 genome editing

Standard methods were used to design and generate desired genomic changes (ARRIBERE *et al*. 2014; KIM *et al*. 2014; PAIX *et al*. 2014; PAIX *et al*. 2015; FAY *et al*. 2021; GHANTA *et al*. 2021).

Oligos including sgRNAs, repair templates, and amplification primers used in this study are listed in Supplementary File 2. CATP-1–fluorescent protein fusion reporters were generated using methods from Dickinson and colleagues (DICKINSON *et al*. 2015; DICKINSON AND GOLDSTEIN 2016). All CRISPR-generated mutations were also confirmed by sequencing.

### Image acquisition and analysis

Confocal fluorescence images (Figure 3) were acquired using an Olympus IX83 inverted microscope with a Yokogawa spinning-disc confocal head (CSU-W1). z-Stack images with a step size of 0.2 µm were acquired using a 100×, 1.35 N.A. silicone oil objective. cellSense 3.1 software (Olympus Corporation) was used for image acquisition. DIC images and fluorescence images (Figures 1 and 3) were obtained using a Nikon Eclipse epifluorescence microscope using 10×, 0.25 N.A. and 40×, 0.75 N.A. objectives. Image acquisition was controlled by Openlab 5.0.2 software (Agilent Inc.). Before they were imaged, animals were mounted on 3% agarose pads and anesthetized using 1 mM levamisole in M9 buffer. Image processing and analysis were carried out using FIJI software (SCHINDELIN *et al*. 2012).

**Figure 1.**
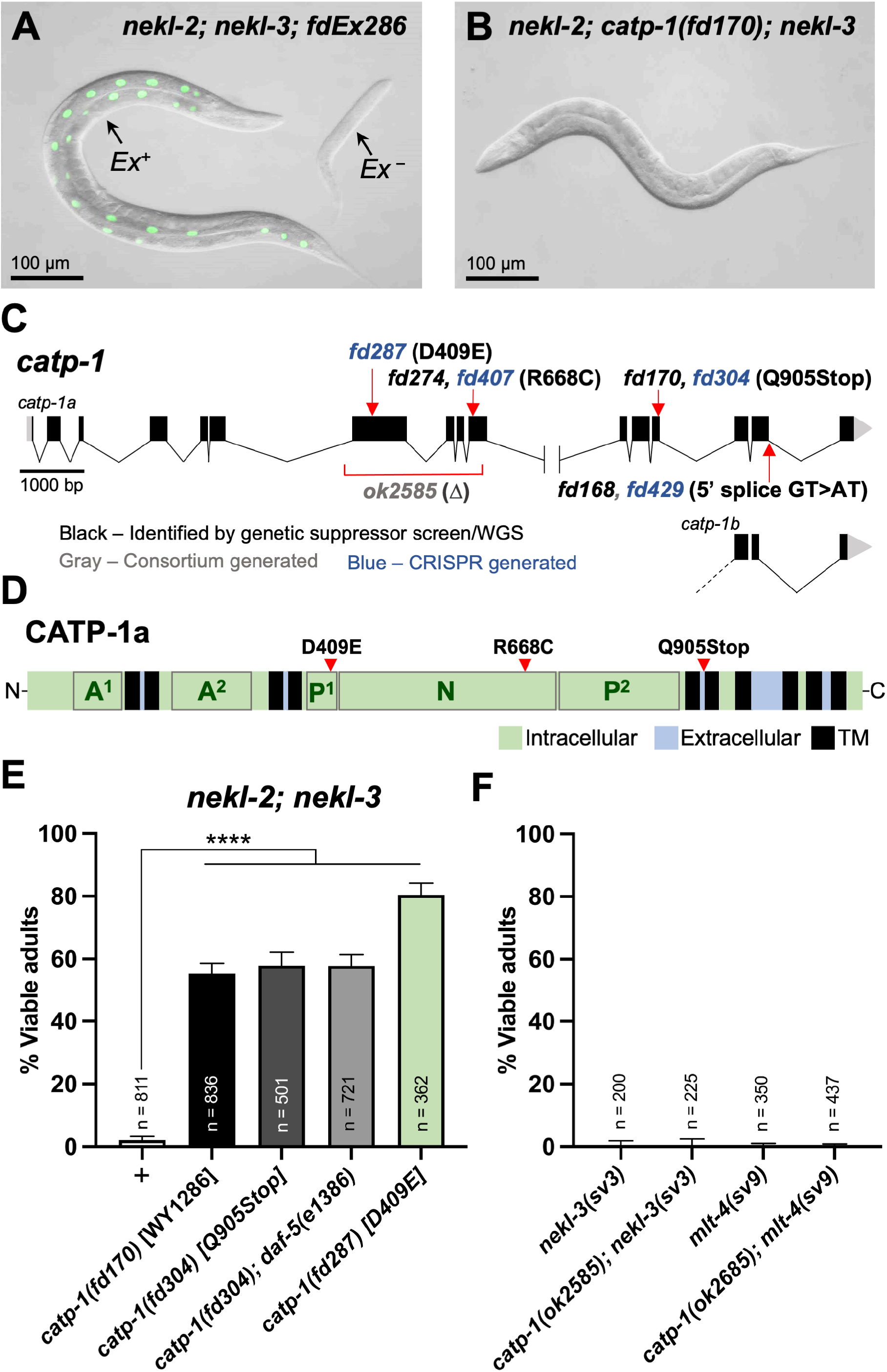
Mutations in *catp-1* lead to suppression of *nekl* molting defects. (A, B) DIC and GFP image overlays of (A) *nekl-2(fd81); nekl-3(gk894345)* and (B) *nekl-2(fd81); catp-1(fd170); nekl-3(gk894345)* strains. Whereas the GFP-positive adult in A carries the *fdEx286* extrachromosomal array (*Ex^+^*) containing rescuing wild-type *nekl-3* genomic sequences and the *sur-5*::GFP reporter, the non-rescued GFP-negative sibling larva (*Ex^−^*) is terminally arrested at the L2/L3 molt. In contrast, ∼50% of *nekl-2(fd81) catp-1(fd170); nekl-3(gk894345)* worms (B) do not exhibit molting defects and reach adulthood. (C) Map of the *catp-1* genomic locus showing the locations of the relevant mutants generated for this study. Alternatively spliced forms of *catp-1a* and *catp-1b*, which differ at C-terminal exons 15 and 16 only, are indicated. (D) Protein diagram of CATP-1a indicating the predicted locations of transmembrane domains (TM); intracellular and extracellular regions; and protein domains including actuator (A^1^, A^2^), phosphorylation (P^1^, P^2^), and ATP-binding (N) domains. The locations of several key amino-acid variants are also indicated. (E) Percentage of viable adults for the indicated genotypes, which include *nekl-2(fd81); nekl-3(gk894345)* mutations in addition to the indicated mutations in *catp-1*; plus sign (+) indicates control strain containing a wild-type *catp-1* allele. (F) Percentage of viable adults for the indicated genotypes. Error bars in E and F indicate 95% confidence intervals; p-values were calculated using Fishers’ exact text; ****p < 0.0001.

### Synchronization and analysis of molting

Assays for molting arrest and genetic suppression were carried out as described (LAZETIC AND FAY 2017a; JOSEPH *et al*. 2018). To generate synchronized populations of L1 larvae, gravid adults were subjected to treatment with bleach, after which eggs were washed at least five times and allowed to hatch in M9 buffer overnight at room temperature with gentle rotation (PORTA-DE-LA-RIVA *et al*. 2012). Hatched larvae were transferred to NGM/OP50 plates, and pharyngeal pumping or P*mlt-10*::GFP–PEST (FRAND *et al*. 2005) expression was recorded every hour at 20°C using an Olympus MVX10 MacroView microscope equipped with a 2× objective. Molting was defined as when ≥50% animals (n = 10) were pumping or when >50% expressed P*mlt-10*::GFP– PEST.

### Statistical analysis and data availability

All statistical tests were performed using GraphPad Prism 10 following established protocols (FAY AND GEROW 2013). Data from all experiments are contained in Supplementary File 3.

## RESULTS

### Loss of the cation exchanger CATP-1 suppresses molting defects in *nekl-2; nekl-3* mutants

In a genetic screen for suppressors of *nekl-2(fd81); nekl-3(gk894345)* synthetic lethality, we isolated strain WY1286 [*nekl-2(fd81) fd170; nekl-3(gk894345)*] (JOSEPH *et al*. 2018). Whereas 2% of *nekl-2(fd81); nekl-3(gk894345)* (hereafter *nekl-2; nekl-3*) mutants developed into viable adults because of molting defects, >50% of *nekl-2 fd170; nekl-3* triple mutants progressed to adulthood within ∼3 or 4 days (Fig 1A, B, E). Backcrossing of WY1286 further suggested that genetic suppression was caused by a single-locus and autosomal recessive mutation. Consistent with this, WGS, in conjunction with the Sibling Subtraction Method (JOSEPH *et al*. 2018), indicated that the suppressor mutation was on chromosome I and identified three point mutations predicted to alter protein coding sequences within the distal right arm of chromosome I. These included a G>A transition resulting in a premature Q905Stop in CATP-1 (CAG>TAG), a C>T transition causing an A341T substitution in F22G12.4 (GCA>ACA), and a G>A transition leading to a G253E substitution in TAF-1 (GGA>GAA). Introduction of an equivalent *catp-1* Q905Stop mutation into *nekl-2; nekl-3* mutants using CRISPR/Cas9 methods (*fd304*) (FARBOUD AND MEYER 2015; FARBOUD *et al*. 2019; FAY *et al*. 2021) led to adult viability of 58%, similar to that observed for *nekl-2 fd170; nekl-*3 animals (Figure 1E; Supplementary File 1).

These findings indicate that *catp-1(fd170)* is the causative suppressor mutation in strain WY1286. We note that *catp-1(RNAi)* failed to suppress molting defects in *nekl-2; nekl-3* mutants, possibly due to insufficient knockdown of CATP-1 by this method.

*catp-1* is predicted to encode a P4-type ATPase alpha subunit, which in conjunction with beta and gamma subunits forms a membrane-associated cation pump (RUAUD AND BESSEREAU 2007). In *C. elegans*, CATP-1 is most similar to CATP-2 and CATP-3—both are 41% identical and 61% similar to CATP-1—and CATP-1 is 30% identical and 49% similar to CATP-4. With respect to human family members, CATP-1 is most similar to the alpha type IIC subfamily members ATP1A2 and ATP1A3, which function as ATP-dependent Na^+^/K^+^ exchangers (KUHLBRANDT 2004; RUAUD AND BESSEREAU 2007; FARLEY 2012). Acting at the plasma membrane, members of this subfamily export three Na^+^ ions and internalize two K^+^ ions during each catalytic cycle, thereby producing an inside–outside electrochemical gradient.The predicted domain structure of CATP- 1a (Figures 1D, S1) includes 10 transmembrane (TM) domains, along with bipartite intracellular actuator (A1/A2) and phosphorylation (P1/P2) domains and a contiguous intracellular ATP-binding (N) domain. A second predicted isoform, CATP-1b, has an altered C terminus and contains sequences through the first 8 TM domains (Fig 1C).The Q905Stop mutations (*fd170* and *fd304*) are predicted to remove the C-terminal 216 and 140 amino acids of CATP-1a and CATP-1b isoforms, respectively (Figure 1C, D, S1).

To determine whether elimination of CATP-1 cation-pump activity is relevant to its role in the context of *nekl* suppression, we used CRISPR/Cas9 to modify a highly conserved aspartic acid residue (D409) within the conserved D-K-S/T-G-T sequence of the P1 domain (Figures 1D and S1) (OHTSUBO *et al*. 1990; RUAUD AND BESSEREAU 2007). During the cation exchange process, this aspartic acid is transiently phosphorylated, leading to the formation of a phosphoenzyme intermediate and a change in conformational state (POST AND KUME 1973; MORTH *et al*. 2011; CLAUSEN *et al*. 2017). Notably, CATP-1 D409E robustly suppressed *nekl-2; nekl-3* synthetic lethality (∼80% viable; Figure 1E), indicating that loss of the CATP-1 ion pump activity is likely critical to its mechanism of *nekl* suppression. The greater suppression level observed for CATP- 1(D409E) relative to that induced by the Q905Stop mutation could be due to the partial activity of the truncated protein produced by the CATP-1(Q905Stop) mutation. Taken together, our results indicate that loss of CATP-1 cation exchange function leads to the suppression of *nekl-2; nekl-3* molting defects.

We also tested whether loss of CATP-1 function could suppress molting defects in stronger loss-of-function alleles of *nekl* or *mlt* genes (YOCHEM *et al*. 2015; LAZETIC AND FAY 2017a). In contrast to *nekl-2; nekl-3* double mutants, *catp-1(ok2585)*, which contains a ∼2,200 bp deletion spanning intron six to exon 10, was unable to suppress the fully penetrant L2/L3 arrest phenotypes of either *nekl-3(sv3)* or *mlt-4(sv9)* single mutants (Figure 1F). These results are similar to our previous observations that loss of the ADM-2 metalloproteinase or induction of the dauer pathway confers only limited suppression of *nekls* (BINTI *et al*. 2022; JOSEPH *et al*. 2022). In contrast, mutations in AP2-associated factors can suppress stronger *nekl* and *mlt* loss-of-function alleles, including molecular nulls (JOSEPH *et al*. 2020).

### *catp-1* mutations identified in two other sequenced strains are insufficient for suppression

WGS of two other suppressor strains, WY1768 (*nekl-2; fd168; nekl-3*) and WY1744 (*nekl-2 fd274; nekl-3*), identified two additional mutations predicted to alter CATP-1. Whereas strain WY1744 exhibited 34% adult viability, strain WY1768 was ∼95% viable, placing it among the strongest suppressors identified by our screen (Figures 1C, 1D, S2A, S2B). Both strains were sequenced after minimal backcrossing and without the Sibling Subtraction Method. Whereas WGS of WY1744 identified nine coding mutations, seven of which were on chromosome I, sequencing of WY1768 identified 14 coding mutations, of which only three were on chromosome I (Figure S2B).

In the case of strain WY1744, the mutation in *catp-1* led to an R668C substitution in a residue directly adjacent to a highly conserved arginine required for ATPase activity (R669; Figure 1C, D, S1) (JACOBSEN *et al*. 2002; RUAUD AND BESSEREAU 2007; MORTH *et al*. 2011; CLAUSEN *et al*. 2017).

Nevertheless, BLAST searches indicate that R668 is less highly conserved than R669 within nematodes; the equivalent position in *Caenorhabditis bovis* and *Wuchereria. bancrofti* CATP-1 is a cysteine and a lysine, respectively. The mutation in strain WY1768 alters the conserved 5’ splice site of *catp-1a* intron 15 (GT>AT; Figure 1C). Failure to incorporate exon 16 would remove the 62 C-terminal amino acids of CATP-1a, including portions of the ninth TM domain, the tenth TM domain, and the C-terminal 21 amino acids (Figures 1C, S1). In contrast, the shorter CATP-1b isoform would not be expected to be altered by the splice-site mutation (Figure 1C).

CRISPR methods were used to introduce analogous R668C (*fd407*) and *catp-1a* intron-15 splice-site (*fd429*) mutations into the wild-type *catp-1* locus in *nekl-2; nekl-3* mutants (Figure 1C).

Surprisingly, neither mutation conferred robust suppression, although *fd407* gave rise to ∼6% adult viability, which was slightly above background (p < 0.001; Figure S2A). These results demonstrate that neither mutation in *catp-1* is sufficient to suppress molting defects in *nekl-2; nekl-3* mutants and implicate other loci as at least partially required for the observed genetic suppression in these strains.

To determine whether the detected mutations in *catp-1* might partially contribute to the suppression observed in WY1744 and WY1768 strains, we attempted to revert the *catp-1* mutations in these backgrounds using CRISPR/Cas9 but were unsuccessful in obtaining the necessary mutations. We also attempted transgenic rescue experiments using *catp-1a* cDNA under the control of a hypodermal-specific promoter (Y37A1B.5; Phyp7::CATP-1), including a construct that encodes an in-frame N-terminal GFP (Phyp7::GFP::CATP-1). Unfortunately, both *catp-1* expression constructs proved to be extremely toxic to worms, even when these constructs were highly diluted with several different co-injection markers. As such, we were unable to determine the extent to which these mutations in *catp-1* might contribute to the suppression detected in these strains. Nevertheless, our findings highlight the importance of follow-up tests to determine the functional role of mutations discovered by WGS, including mutations in genes that have been shown to be sufficient to produce the relevant phenotype.

### CATP-1 is expressed in apical puncta in the epidermis

CATP-1 was previously shown to be expressed and to function in epidermal cells including the hypodermal syncytia in the head (hyp1 to hyp6), midbody (hyp7), and tail (hyp8 to hyp11) (RUAUD AND BESSEREAU 2007). hyp7 is also the tissue focus for NEKL and MLT activities and is the major contributor to the larval and adult cuticles (YOCHEM *et al*. 2015; LAZETIC AND FAY 2017a; LAZETIC AND FAY 2017b). CATP-1 was also reported to function in glial cells that surround touch-sensory neurons in the head (JOHNSON *et al*. 2020). In addition, modENCODE RNA sequencing data available on WormBase indicate that CATP-1 is expressed throughout development, including in embryos and all four larval stages, with highest levels corresponding to dauer larvae (LI *et al*. 2014; HARRIS *et al*. 2020).

To further examine the tissue expression and subcellular localization of CATP-1, we obtained endogenous CRISPR-tagged variants in which the CATP-1a isoform was tagged at its C terminus with either GFP or mScarlet. Consistent with previous studies, we observed clear expression of CATP-1::GFP and CATP-1::mScarlet in the hyp7 midbody, as well as in head and tail epidermal cells young adults (Figure 2A–G); expression at earlier larval stages was more difficult to analyze because of competing auto-fluorescence coming from the intestine. With respect to subcellular localization, we observed faint CATP-1::GFP in punctate and reticular structures in the apical region of hyp7 underlying the cuticle, which may correspond to membrane or juxtamembrane accumulations including clathrin-coated pits and/or recycling endosomes (Figure 2A–C).

**Figure 2.**
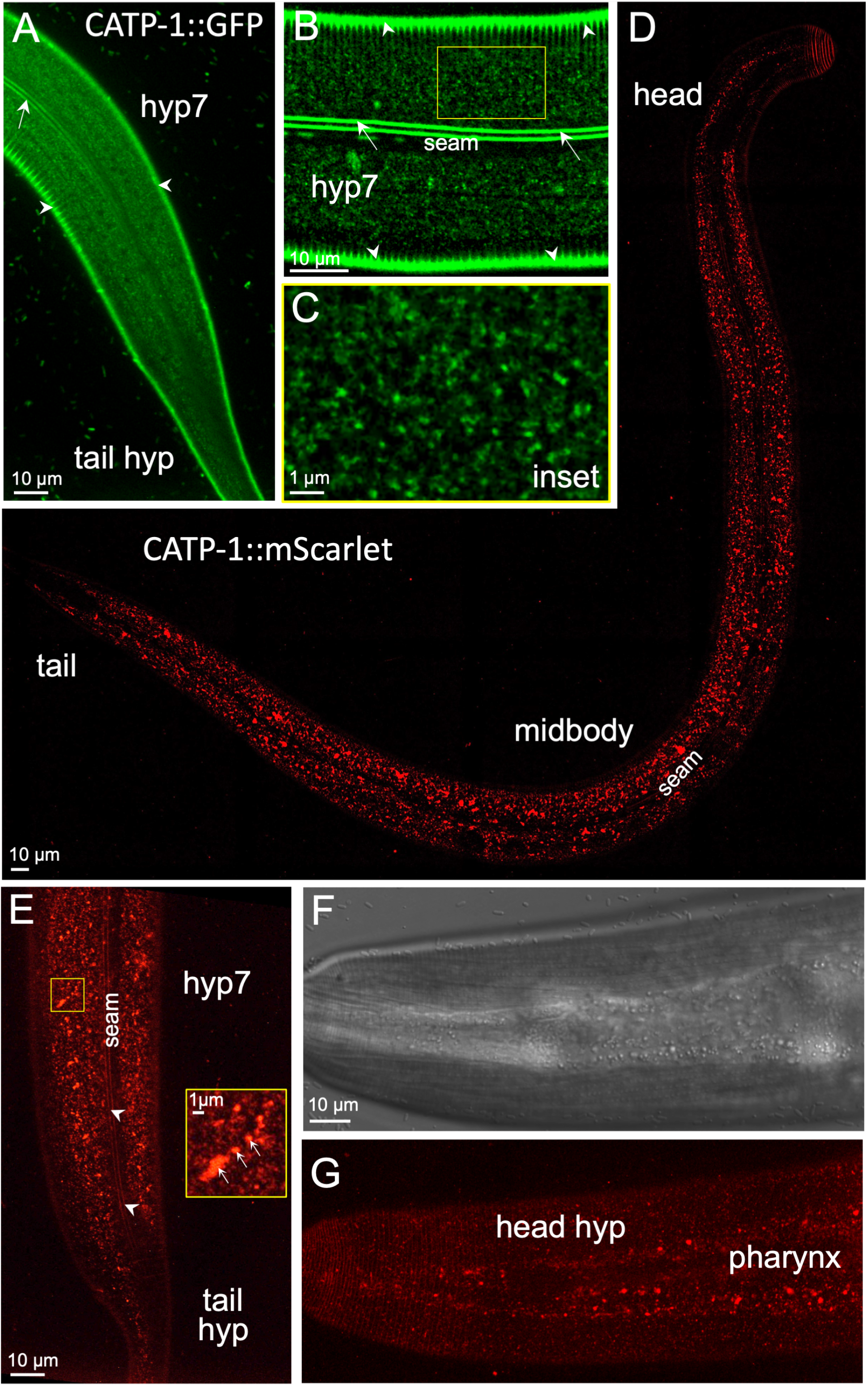
CATP-1 is expressed predominantly in punctate structures within the epidermis. (A–G) Fluorescence (A–E, G) and DIC (F) images of worms expressing endogenously tagged CATP-1::GFP (A–C) and CATP-1::mScarlet (D–G), in which sequences encoding fluorescent proteins were integrated at the C terminus of the *catp-1a* isoform. All panels show young adults. (A) Apical regions corresponding to the midbody (hyp7 syncytium) and tail (hyp8–11) are indicated. (B) Midbody apical section of hyp7 and (C) enlarged inset showing punctate and reticular structures. In A–C autofluorescence of the dorso-ventral cuticle (white arrowheads) and lateral cuticular alae (white arrows) are indicated, along with the location of the seam cell (panel B), in which CATP-1::GFP was not detected. (D) Whole-body apical image of a young adult expressing CATP-1::mScarlet with relevant regions indicated. (E) CATP-1::mScarlet expression in the apical midbody and tail regions along with enlarged inset. Note the presence of fluorescence signals in variably sized punctate structures including accumulation of mScarlet in larger structures that likely correspond to late endosomes and lysosomes (white arrows in inset). White arrowheads in panel E indicate autofluorescence of cuticular alae. (F) DIC and (G) corresponding fluorescence images of CATP-1::mScarlet in the apical head region showing expression in both the anterior hypodermis (hyp1–6) and pharynx, in which larger puncta accumulate.

Likewise, faint CATP-1::mScarlet fluorescence was observed in apical epidermal puncta but was also detected at higher levels in more medial compartments, which may correspond to late endosomes and lysosomes (Figure 2D–E). This difference in GFP and mScarlet localization is likely explained by the observation that red fluorescent proteins can be cleaved from their translational fusion partners leading to their accumulation in acidic compartments of the endo- lysosome system in worms (CLANCY *et al*. 2023). In addition, CATP-1::mScarlet was observed in punctate structures within the region of the pharynx (Figure 2F, G). In contrast, both CATP- 1::GFP and CATP-1::mScarlet were largely absent from epidermal seam cells (Figure 2A, B, D, E). The absence of apparent CATP-1 expression in nervous system–associated structures, such as head glia, may reflect sensitivity limitations of our reporters. Collectively, our data are consistent with CATP-1 playing a role in the epidermis during the molting process.

### Suppression of *nekl* molting defects by *catp-1* loss does not occur via a dauer-induction mechanism

Mutations in *catp-1* suppress larval arrest after exposure to DMPP (RUAUD AND BESSEREAU 2006; RUAUD AND BESSEREAU 2007). Suppression of DMPP toxicity was also observed under environmental conditions that induce the formation of dauers and in genetic mutants that cause constitutive dauer entry (RUAUD AND BESSEREAU 2006; RUAUD *et al*. 2011). Data further suggested that suppression of DMPP might correlate with entry into L2d, a pre-dauer stage from which worms can either transition into dauers (L3d) or resume continuous development by entering L3. Notably, the duration of L2d is typically ∼1.2–2-fold longer than that of L2, depending on the strength and nature of the inducing signal, and suppression of DMPP toxicity by *daf-2* mutants is positively correlated with the extent of L2 elongation (GOLDEN AND RIDDLE 1984; RUAUD *et al*. 2011; KARP 2018; BINTI *et al*. 2022). Consistent with this, loss of *catp-1* was reported to extend the length of L2 by ∼1.6-fold but to have no effect on the duration of L1 or other larval stages (RUAUD AND BESSEREAU 2007).

Recently, we showed that induction of L2d is sufficient to suppress molting defects in ∼50% of *nekl-2; nekl-3* mutants but is much less effective at suppressing defects in stronger loss-of-function alleles of *nekls* or *mlts*, a suppression profile similar to that of *catp-1* (Figure 1F) (BINTI *et al*. 2022). We thus hypothesized that loss of *catp-1* may suppress *nekl-2; nekl-3* molting defects by inducing L2d or an L2d-like state. As a first test, we introduced a dauer-defective allele of *daf-5* (*e1386*) (RIDDLE *et al*. 1981) into *nekl-2 catp-1; nekl-3* mutants to inhibit entry into L2d. Notably, we had previously shown that *daf-5(e1386)* significantly reduces the ability of dauer-inducing conditions to suppress *nekl-2; nekl-3* molting defects (BINTI *et al*. 2022).

However, we observed no difference in the level of suppression in *nekl-2 catp-1; daf-5; nekl-3* mutants versus *nekl-2 catp-1; nekl-3* mutants (Figure 1E), suggesting that loss of *catp-1* might not suppress *nekl* molting defects through an L2d mechanism or that loss of *catp-1* induces L2d via a mechanism that is independent of DAF-5 function.

To further test for the induction of L2d in *catp-1* mutants, we examined the timing of larval molts in wild type and *catp-1* mutants using two different methods. In one assay, we followed molts by scoring pharyngeal pumping in a synchronized population of worms released from early L1-larval arrest; loss of pumping occurs when animals enter the lethargus phase of the molting cycle and is commonly used to measure the duration of larval-stages (SINGH AND SULSTON 1978; LAZETIC AND FAY 2017b). Using two different strong loss-of-function alleles of *catp-1* (*ok2585* and *kr17*), we observed a slight lengthening of both L1 and L2 by ∼1.1- to 1.2-fold in *catp-1* mutants but no strong or specific effect at the L2 stage (Figure 3A, C). As a further assessment, we similarly timed molting cycles using the P*mlt-10*::GFP–PEST marker, which peaks in expression for several hours during each molting cycle (FRAND *et al*. 2005). Based on this marker, *catp-1* mutants displayed an ∼1.2-fold increase in the length of L1 but exhibited little or no increase in the length of L2 (Figure 3B, C). As such, we failed to detect a specific lengthening of the L2 stage in *catp-1* mutants, nor were strong effects on stage length observed at L3 or L4 stages (Figure 3B, C).

**Figure 3.**
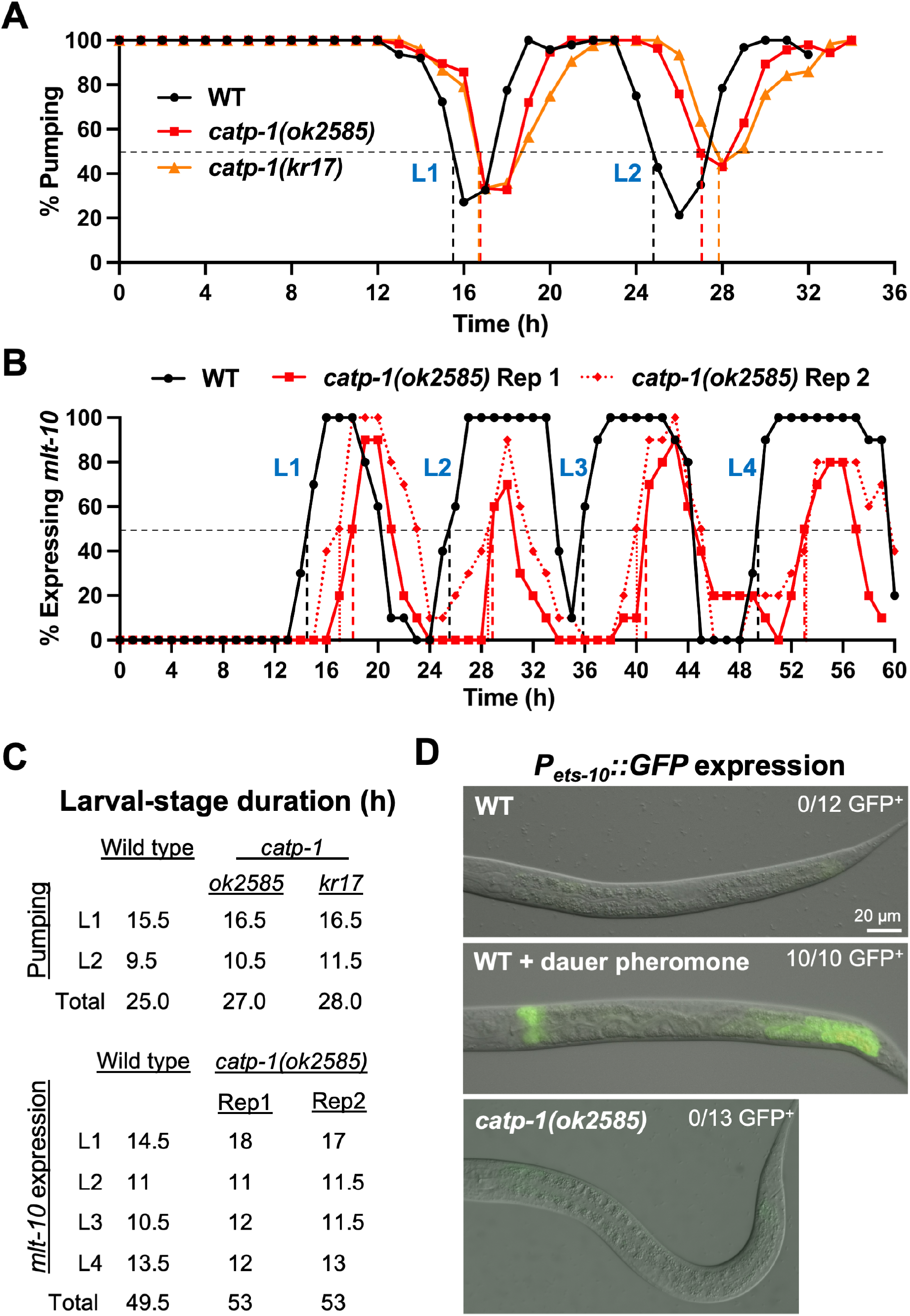
Loss of *catp-1* does not substantially extend the length of the L2 stage nor does it induce an L2d marker. (A) Cessation of pharyngeal pumping was used to measure the duration of L1 and L2 stages in the indicated backgrounds. (B) P*mlt-10*::GFP–PEST expression was used to measure the lengths of larval stages in wild-type and *catp-1* larvae; two replicates are shown for *catp-1*. Dashed vertical lines in A and B indicate the point at which ≥50% of larvae had entered lethargus for a given stage. (C) Quantification of larval-stage lengths for A and B. (D) Expression of P*mlt-10*::GFP–PEST in L2 larvae under the indicated conditions. In contrast to wild type with dauer pheromone, neither untreated wild type nor *catp-1* mutants express the L2d marker.

As a final test for L2d induction by loss of *catp-1*, we determined whether an established marker for L2d is inappropriately expressed under normal growth conditions in *catp-1* mutants. P*ets-10*::GFP is mostly undetectable in fed wild-type larvae but is expressed robustly after wild-type larvae are treated with dauer-inducing pheromones (Figure 3D) (SHIH *et al*. 2019; BINTI *et al*. 2022). Notably, *catp-1* mutant larvae did not express the L2d marker under normal growth conditions. Taken together, our data failed to support the model that loss of CATP-1 suppresses molting defects in *nekl-2; nekl-3* mutants by inducing L2d.

## DISCUSSION

Here we show that loss of the CATP-1 Na^+^/K^+^ exchanger led to the suppression of molting defects in *nekl-2; nekl-3* mutants. We also show that a mutation designed to specifically abolish CATP-1 ion-pump activity was sufficient to cause suppression. This is consistent with a recent report implicating CATP-1 pump activity in glia (JOHNSON *et al*. 2020) but contrasts with data from Ruaud and colleagues suggesting pump-independent functions for CATP-1 within the context of DMPP-toxicity suppression (RUAUD AND BESSEREAU 2007; JOHNSON *et al*. 2020). These differences may be due to distinct functional requirements for CATP-1, because of technical differences in expression methods, or because of possible independent effects of the two CATP- 1 isoforms. Consistent with findings by Ruaud and colleagues, we observed expression of CATP- 1 throughout the epidermis (RUAUD AND BESSEREAU 2007). Moreover, we observed CATP-1 in punctate and reticular structures at or near the apical surface of epidermal cells, suggesting that CATP-1 may function at the plasma membrane and possibly endosomes, broadly consistent with previous studies on P4-type Na^+^/K^+^ pumps (KAPLAN 2002; CLAUSEN *et al*. 2017) (and see below). Collectively, our data indicate that *nekl* molting defects may be partially alleviated by altering the concentration or distribution of intracellular solutes or by changes in electrochemical gradients.

Based on prior studies of *catp-1* and other DMPP-toxicity suppressors (RUAUD AND BESSEREAU 2006; RUAUD AND BESSEREAU 2007; RUAUD *et al*. 2011), we tested the appealing hypothesis that loss of CATP-1 leads to *nekl* suppression by inducing a pre-dauer L2d state, which we recently showed can bypass L2/L3 molting defects in *nekl-2; nekl-3* mutants (BINTI *et al*. 2022). In contrast to expectations, however, our evidence did not support this model. Moreover, we were unable to recapitulate a previous finding that loss of *catp-1* leads to a marked lengthening of the L2 stage, a hallmark of L2d (RUAUD AND BESSEREAU 2007). Although we have no simple explanation to account for these differences, we note that disparities could potentially arise from altered growth conditions or uncharacterized mutations present in strain backgrounds. It is also worth noting that *catp-1* was suggested to suppress DMPP toxicity through a dauer-independent mechanism (RUAUD AND BESSEREAU 2007). Correspondingly, although the suppression of DMPP toxicity by other dauer-inducing mutations correlated strongly with increased L2 duration, it did not correlate with the ability of these mutants to induce mature dauer larvae (RUAUD AND BESSEREAU 2007; RUAUD *et al*. 2011). Thus, while not fully resolved, our collective data appear to suggest roles for CATP-1 that are independent of the dauer pathway and L2d induction.

WGS of *nekl* suppressor strains identified three isolates containing unique mutations that alter the coding region of *catp-1* [WY1286 (*fd170*), WY1768 (*fd168*), and WY1744 (*fd274*)].

Traditionally, the detection of variants at the same locus in two or more independent isolates is thought to imply that the common locus is likely responsible for causing the phenotype of interest. This logic is particularly compelling if mutations at that locus have already been shown to be causative and sufficient, as was the case for *catp-1* (WY1286). We therefore anticipated that the mutations impacting the coding region of *catp-1* in strains WY1768 and WY1744 would be responsible for the observed suppression. Surprisingly, CRISPR phenocopy of these *catp-1* variants failed to suppress *nekl-2; nekl-3* molting defects, indicating that they are insufficient for suppression. Unfortunately, we were unable to determine the extent to which these mutations might partially contribute to the observed suppression.

It was previously shown that the CATP-1a isoform is sufficient for rescuing both DMPP toxicity (RUAUD AND BESSEREAU 2007) and neuronal sensory phenotypes in *catp-1* mutants (JOHNSON *et al*. 2020), indicating that CATP-1a may be functionally sufficient in epidermal and glial cells, respectively. Interestingly, the failure of the GT>AT *catp-1a* intron 15 splice-site mutation to suppress *nekl-2; nekl-3* mutants suggests that the CATP-1b isoform may be able to partially compensate for the loss of CATP-1a. We note that an alternative in-frame splice donor sequence (GT) is not present within intronic sequences immediately downstream of the splice-site mutation, and although several upstream GT sequences in exon 15 could potentially serve this role, they would lead to the partial removal of the CATP-1a ninth TM domain. Notably, CATP-1b is not predicted to encode a ninth or tenth TM domain, and functional P4-ATPases have been reported to contain as few as six TM domains, with TM1–6 making up the functional core (KUHLBRANDT 2004; MORTH *et al*. 2011). Taken together, our data suggest that CATP-1a and CATP-1b may be partially functionally redundant and that both isoforms may encode functional Na^+^/K^+^ pumps.

In prior studies we have shown that NEKL-2 and NEKL-3 localize to several endosomal trafficking compartments, that depletion of NEKL-2 and NEKL-3 leads to morphologically and functionally altered trafficking compartments, and that mutations in several membrane trafficking factors can suppress molting defects associated with *nekl* loss (YOCHEM *et al*. 2015; LAZETIC AND FAY 2017a; LAZETIC *et al*. 2018; JOSEPH *et al*. 2020; JOSEPH *et al*. 2023). Thus, it is possible that loss of *catp-1* may lead to a partial suppression of *nekl* molting defects through direct or indirect effects on membrane trafficking. Some support for this model comes from published studies linking cation levels and electrochemical gradients to effects on membrane trafficking. For example, depletion of intracellular potassium leads to the inhibition of clathrin-coated pit formation, potentially by preventing clathrin from interacting with its adapter proteins (LARKIN *et al*. 1983; HEUSER AND ANDERSON 1989; HANSEN *et al*. 1993). Correspondingly, we found that inhibition of AP2 clathrin-adapter activation can suppress *nekl* molting and trafficking-associated defects (JOSEPH *et al*. 2020). Na^+^/K^+^ pumps also affect the acidification and function of early endosomes by impacting the movement of H^+^ across endosomal membranes, and the pharmacological inhibition of Na^+^/K^+^ pumps leads to endosomal recycling defects (FUCHS *et al*. 1989; ROSEN *et al*. 2004; ROSEN *et al*. 2006; FELDMANN *et al*. 2007). Finally, ionic gradients have been shown to affect vesicle morphology and motility through changes in membrane tension and altered cytoskeletal dynamics (NUNES *et al*. 2015; SIMUNOVIC AND VOTH 2015; MURRAY *et al*. 2017; MERCIER *et al*. 2020; SARIC AND FREEMAN 2021).

In addition to the above-mentioned effects on membrane trafficking, Na^+^/K^+^ pumps affect various cell signaling pathways, which could represent another means by which *catp-1* loss may impact NEKL functions and molting (DAVIS *et al*. 1995; MOHAMMADI *et al*. 2001; LIANG *et al*. 2006; DOI AND IWASAKI 2008; MATCHKOV AND KRIVOI 2016; CLAUSEN *et al*. 2017; PIVOVAROV *et al*. 2018).

Likewise, analogous to its function in glia, CATP-1 could potentially modify the activity of neurons that are closely associated with the epidermis, which might conceivably impact the molting process (JOHNSON *et al*. 2020). At present, however, it remains to be determined precisely how *catp-1* and changes in cation concentration can compensate for *nekl* loss of function. Future studies will examine how mutations in *catp-1* and other identified *nekl* suppressors impact membrane trafficking processes and whether they suppress one or more membrane trafficking defects associated with *nekl* loss of function.

## ACKNOWLEDGMENTS

We thank Amy Fluet for editing this manuscript,Laura Bianchi (University of Miami) for reagents and helpful discussions, Adison Linder, for experimental support, and the *C. elegans* Gene Knockout Consortium. This work was supported by R35 GM136236 and by an Institutional Development Award (IDeA) from the National Institute of General Medical Sciences of the National Institutes of Health under (2P20GM103432).

## Notes

### Competing Interest Statement

The authors have declared no competing interest.

### Summary of Updates

We are correcting some typos that were present in the supplementary files of the first submission.

