## Supplementary figures and images for "Loss of the Na^+^/K^+^ cation pump CATP-1 suppresses *nekl*-associated molting defects"

### Supplementary Figure 1

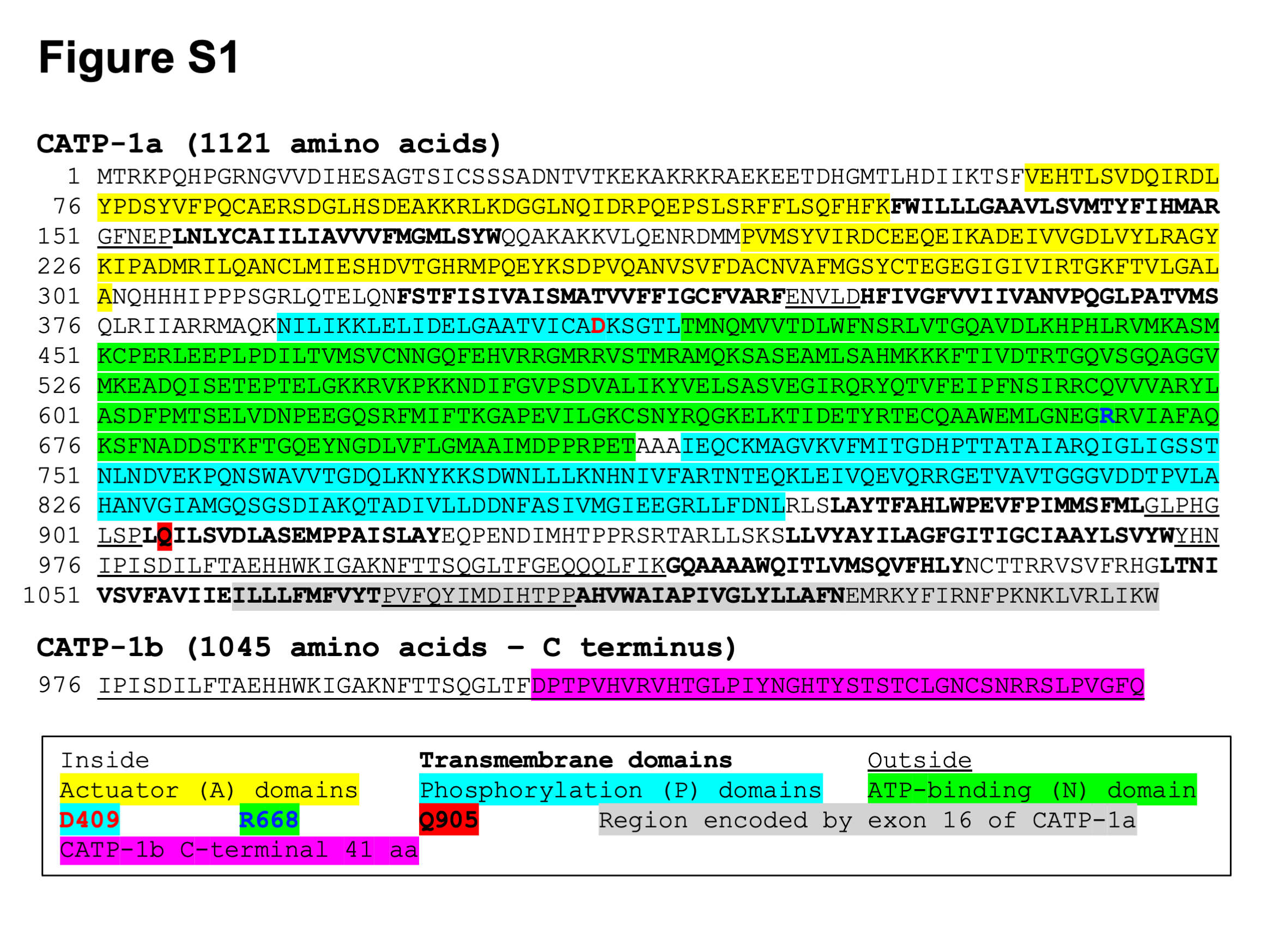

### Supplementary Figure 2

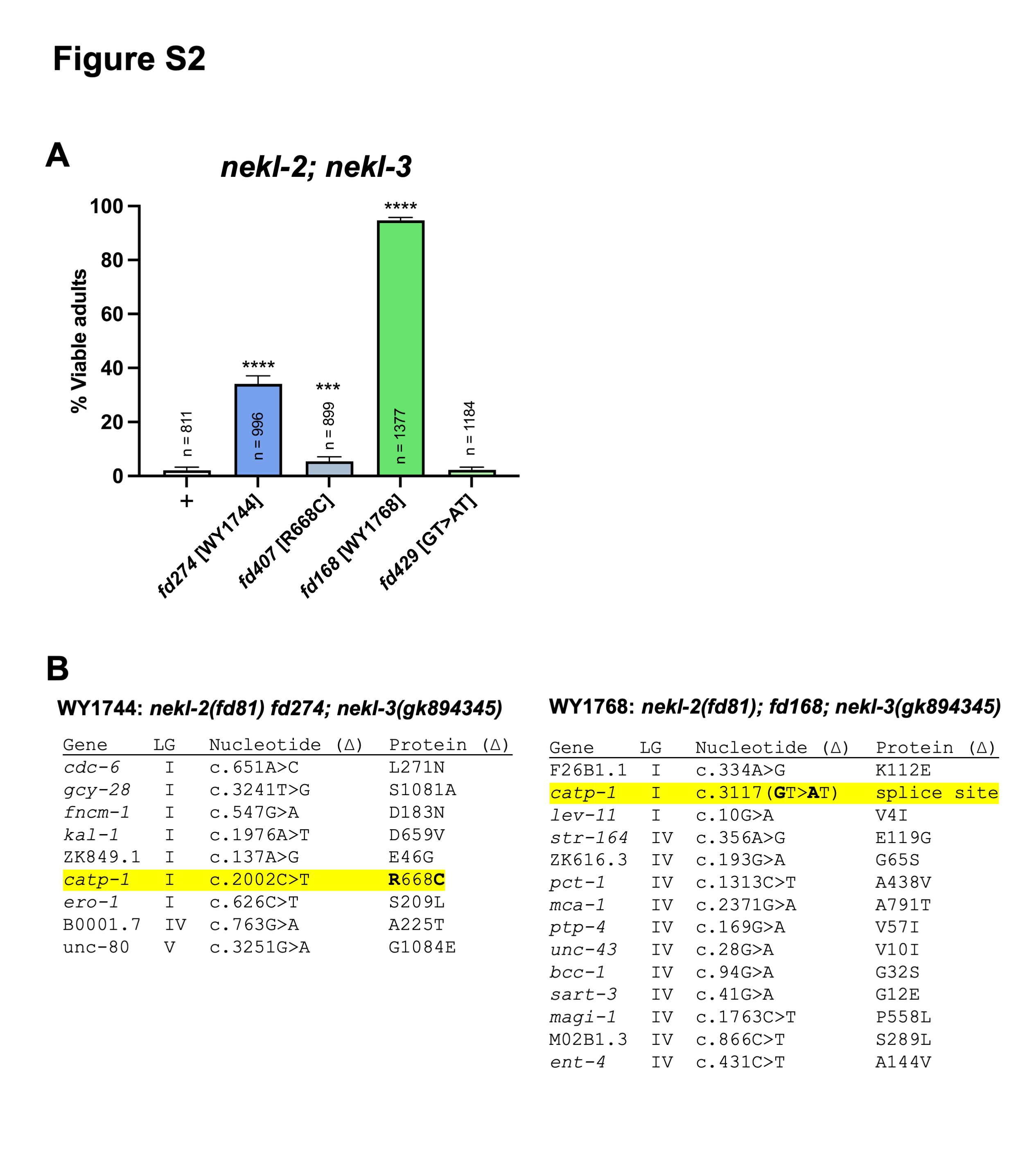
